# Beyond the phloem: A general role of the Arabidopsis *OCTOPUS* gene family in controlling plant growth vigour

**DOI:** 10.1101/2024.05.17.594124

**Authors:** Simona Crivelli, Kai Bartusch, M. Aguila Ruiz-Sola, Mario Coiro, Signe Schmidt Kjølner Hansen, Elisabeth Truernit

## Abstract

*OCTOPUS* (*OPS*) and *OCTOPUS-LIKE 2* (*OPL2*), two homologous genes, were previously identified as important regulators of phloem differentiation in Arabidopsis roots, impacting root growth when their function is lost. Here, we investigated the roles of the other three *OPS* homologs in Arabidopsis, *OPL1*, *OPL3*, and *OPL4*. We employed promoter activity analyses, protein localisation studies, functional complementation assays, analysis of single and multiple mutant combinations, and growth assays, including exposure to CLE45 and brassinosteroid pathway modulators. The *OPS/OPL* genes exhibit overlapping expression patterns and functions. Multiple mutant combinations revealed a high degree of redundancy, with *OPS* and the phloem domain playing a major role in controlling plant growth. While phloem phenotypes are not exacerbated in higher-order mutants, plant growth vigour is nevertheless more severely impacted than in *ops opl2*. Our results suggest a novel role of the *OPS/OPL* genes in broadly controlling plant growth and development, potentially through the modulation of meristematic activity via brassinosteroid pathways.

## Introduction

The Arabidopsis root serves as an ideal system for studying the fundamental principles of plant organ growth. Within the root meristem, located at the tip of the root, stem cells are in contact with a small number of quiescent centre (QC) cells. The stem cells divide and give rise to various cell types which are organised in layers (files) around the core of the root, the stele (Dolan et al., 1993). The stele houses tissues responsible for transporting water and carbohydrates, namely xylem and phloem. Each of the two phloem poles in the root contains one protophloem and one metaphloem sieve tube cell flanked by two companion cells (Mähönen et al., 2000).

Root cell file growth involves a delicate interplay between cell division in the root meristem, elongation, and differentiation processes along the longitudinal axis of the root. Furthermore, each tissue type follows its own developmental timing, making coordination between their growth crucial. For instance, among all cell files, protophloem cells cease dividing, commence elongating, and then differentiate closest to the root meristem (Lavrekha *et al*., 2017). Given that the protophloem is crucial for unloading solutes into the root tip (Ross-Elliott *et al*., 2017), this early differentiation probably guarantees sufficient energy supply to sustain meristem activity.

Brassinosteroids (BRs) are a group of plant hormones crucial for regulating root cell division and elongation in a concentration-dependent manner (Vukašinović *et al*., 2021). Briefly, perception of BRs by the BRASSINOSTEROID INSENSITIVE 1 (BRI1) receptor kinase or its two homologs, BRI1-LIKE 1 (BRL1) and BRL3, leads to the degradation of BRASSINOSTEROID-INSENSITIVE 2 (BIN2), a GLYCOGEN SYNTHASE KINASE 3 (GSK3)-like kinase. Subsequently, the transcription factors BRI1-EMS-SUPPRESSOR 1 (BES1) and BRASSINAZOLE-RESISTANT 1 (BZR1) activate the transcription of BR-induced genes. In the absence of BRs, BIN2 translocates to the nucleus and phosphorylates BES1 and BZR1, resulting in the transcriptional repression of BR-induced genes (Nolan *et al*., 2020).

Expression of *BRI1*, which is usually ubiquitously expressed (Friedrichsen *et al*., 2000), in the protophloem of the triple BR receptor mutant *bri1 brl1 brl3* is sufficient to restore the severely impaired root growth in this mutant (Kang *et al*., 2017; Graeff *et al*., 2020). Thus, BR signalling in the phloem appears to be crucial for proper root growth. Importantly, this also highlights the essential role of the phloem as a central organiser of root growth beyond its physiological function.

OCTOPUS (OPS), a highly disordered membrane associated protein with no described functional domains, is specifically expressed in the developing protophloem and adjacent metaphloem cell files, where it was shown to sequester BIN2 to the plasma membrane, thus promoting BR signalling (Truernit *et al*., 2012; Anne *et al*., 2015). In addition, OPS interferes, likely directly, with the sensing of the phloem expressed CLAVATA3/EMBRYO SURROUNDING REGION-RELATED 33 (CLE33) and CLE45 peptides by their cognate receptor, BARELY ANY MERISTEM 3 (BAM3), a leucine-rich repeat receptor-like kinase (LRR-RLK) (Breda *et al*., 2019; Carbonnel *et al*., 2023). OPS loss-of-function mutants exhibit interrupted protophloem files with non-differentiated cells and, likely also as a consequence of this, *ops* root growth is adversely affected (Truernit *et al*., 2012). Thus, although its exact molecular function remains elusive, OPS is crucial for promoting the coordinated differentiation of Arabidopsis protophloem sieve element cell files.

In Arabidopsis, *OPS* is part of a small gene family with four additional members: *OCTOPUS-LIKE 1* (*OPL1*), *OPL2*, *OPL3*, and *OPL4*. Within the family, *OPS*, *OPL1*, and *OPL2* are more closely related to each other and form class I, whereas *OPL3* and *OPL4* together form class II (Nagawa *et al*., 2006; Ruiz-Sola *et al*., 2017). Previously, we showed that OPS and OPL2 act redundantly in phloem development. In addition to the protophloem defects, the metaphloem sieve tubes, responsible for long-distance solute transport, also display differentiation defects in *ops opl2*. Root growth is even more affected in *ops opl2*, and the number of cells in the stele is decreased compared to *ops*. In contrast to OPS, however, OPL2 is expressed throughout the root meristem (Ruiz-Sola *et al*., 2017).

Another interesting feature about OPS/OPL proteins is their polar localisation in the cell. In roots, all studied OPS/OPL proteins fused to fluorescent proteins exhibited membrane association directed towards the shootward direction (Truernit *et al*., 2012; Ruiz-Sola *et al*., 2017; Wallner *et al*., 2023). Recently, it was also shown that some OPL proteins are polarly localised in cotyledon stomatal precursor cells, where, after asymmetric division, they segregate into meristemoids, promoting their meristematic activity and delaying guard cell differentiation. In this context, OPL2 seems to play the major role, as multiple *opl* mutants do not display more severe defects than *opl2*. In general, however, the mutant phenotypes in cotyledons were mild, manifesting as slightly increased epidermis cell sizes and reduced cell numbers, while the effects on guard cell development in plants expressing OPL2 gain-of-function versions were comparably stronger (Wallner *et al*., 2023).

With the aim of understanding the contribution of all *OPS/OPL* family genes to phloem development and root growth, in this study we independently generated and carefully investigated multiple *ops/opl* mutant combinations. We demonstrate that, except for *OPS* and *OPL2*, these genes display no redundancy in the control of root phloem sieve tube differentiation. However, they all redundantly contribute to plant growth vigour, most likely by promoting meristematic activity through modulation of BR signalling. Among the *OPS/OPL* genes, *OPS* is the only gene expressed specifically in the phloem, while most of them are expressed more broadly in the root meristem. Nevertheless, all OPL proteins are, at least partially, functionally equivalent to OPS. Our results suggest a scenario in which a higher expression of any *OPS/OPL* gene in the developing phloem files is sufficient to ensure optimal root growth, once again highlighting the central role of the phloem as organiser of plant growth.

## Materials and methods

### Plant material and growth conditions

*Arabidopsis thaliana* (L.) Heynh (ecotype Columbia) served as the wild type throughout the experiments. *ops-2* (SALK_139316) and *opl2-1* (SALK_004773) were described before (Truernit *et al*., 2012; Ruiz-Sola *et al*., 2017). Additionally, mutants SALK_152815, SALK_138940, and SALKseq_072024 (also refer to Fig. S3) were obtained from the SALK collection (Salk Institute, San Diego, CA, USA) (Alonso *et al*., 2003). Higher-order mutants were created via crossing.

If not stated otherwise, for growth experiments, plants were cultivated either in growth chambers on soil (light: 130 µmol m^-2^ s^-1^ for 16 hours, temp: 20°C, humidity: 60%), or on standard media (1/2 Murashige and Skoog (MS) including vitamins (Duchefa Biochemie), 0.5 g/L MES monohydrate, and 1.2% plant agar) under continuous light (110 µmol m^-2^ s^-1^). For CLE45 treatments, seeds were sown directly on media containing the CLE peptide (GenScript Biotech). For treatments with bikinin, brassinolide, or brazzinazole, seedlings were shifted to media containing these components (or the corresponding amount of the solvent DMSO as mock control) after 4 days of growth on standard media. Root growth after transfer was assessed by measuring the change in root length between the time of transfer and three days thereafter.

### Tissue preparation and microscopy

For confocal laser-scanning microscopy of roots, a Zeiss LSM 780 was used. Seedlings were either immersed in ClearSee and stained with Calcofluor White, or, for live imaging, roots were stained with propidium iodide (Kurihara *et al*., 2015; Ursache *et al*., 2018).

Light and fluorescence wide-field microscopy were performed with a Zeiss Axio Imager Z2 or Leica Microscopes M205 and DM2500 using the appropriate filters. Cotyledons for vascular complexity assessment were fixed and cleared before microscopy (Ruiz-Sola *et al*., 2017). For measuring the length of stem cortex cells, parts of fully elongated floral stems were cleared in ClearSee, and images of the cortex cells were taken using wide-field light microscopy. Generation of root cross sections and staining for ß-GLUCURONIDASE (GUS) activity was described before (Ruiz-Sola *et al*., 2017). Cross sections were stained with 0.01% toluidine blue, and only cross sections in which the outer two metaxylem cells were differentiated while the metaxylem cell in the middle was still immature were considered for phloem phenotype analysis. Phloem transport assays were conducted on 8-day-old seedlings following published methods (Serivichyaswat *et al*., 2022). Root tips were inspected for CFDA signal after one hour.

### Molecular biological methods / cloning / plant transformation

Standard molecular biology methods were applied for DNA and RNA extraction, RT-PCR, qPCR, and genotyping. Constructs were generated using the Gateway® Technology (Invitrogen). Cloning strategies and primer sequences can be found in Table 1 and 2. Cloning of *OPS* and *OPL2* reporter constructs was described previously (Ruiz-Sola *et al*., 2017). For Arabidopsis transformation we used a modified floral dip method (Logemann et al., 2006). For all transgenic plants generated, analysis was performed on at least 10 independent T2 lines, with three representative T-DNA single-insert lines chosen for further analysis in homozygous T3 lines. For qPCR, three biological replicates (RNA of a pool of 30 8-day-old seedlings each) were used. The expression of three housekeeping genes (*ACT2*, *PP2AA3*, and *UBI10*) was used as control. As they all showed the same expression pattern, *ACTIN2* was chosen as reference gene.

### Measurements and statistical analyses

For root length analysis, growth plates were scanned. For analysis of root cell parameters, images of root meristems taken with a confocal laser-scanning microscope or of cross sections using wide-field light microscopy were used. For rosette growth parameters, pictures of plants growing on soil were taken with a camera. For all measurements, ImageJ software packages were used. For root meristem size, the length from the quiescent centre to the first cortex cell double the size of the previous one was measured. Root meristem diameter was measured at the end of the meristem. For analysis of relative root growth, data were normalised by dividing the root length of treated plants by the mean root length of the corresponding mock-treated plants (for BR treatments the growth between two timpoints after transfer to the treatment media was used for this calculation).

Statistical analyses were conducted using the RStudio software. Unless stated otherwise, the Shapiro-Wilk normality test for data distribution and the Kruskal-Wallis test followed by Dunn’s multiple comparison post hoc test for non-parametric data was employed. All growth assays were repeated at least once. Cotyledon vascular complexity (number of closed loops and branching points) was assessed based on a published method (Kastanaki *et al*., 2022). Cotyledons were grouped into different categories and statistical significance was calculated by Fisher’s exact test on the groups. qPCR and statistical analysis of gene expression were performed according to standard protocols. All boxplots show the 25^th^ and 75^th^ percentiles and the median, whiskers represent ± 1.5 interquartile range.

## Results

### *OPS and OPL* promoter activity overlaps in several tissues and cell types

To examine the expression of all *OPS*/*OPL* gene family members, we created plants expressing the reporter gene *ß-GLUCURONIDASE* (*GUS*) under the control of the *OPL1*, *OPL3*, and *OPL4* promoters. For a direct comparison of promoter-driven *GUS* expression in seedlings, we also included the previously described *OPS* and *OPL2* GUS-reporter plants in our analysis (Fig. 1) (Ruiz-Sola *et al*., 2017). Both the *OPL3* and *OPL4* promoters were highly active in the RAM, with stronger *OPL4* promoter activity observed around the quiescent centre and in columella cells, while *OPL3* promoter activity was more uniform in the RAM but absent from the columella cells (Fig. 1d,e). Although *OPL2* is also expressed throughout the RAM (Ruiz-Sola *et al*., 2017), its promoter activity was barely detectable in several independent GUS-reporter lines (Fig. 1c), indicating a weaker expression in this tissue compared to *OPL3* and *OPL4*. The *OPL1* GUS-reporter plants showed no expression in the RAM, but clear *OPL1* promoter activity was observed specifically in the developing metaxylem cells in the more mature part of the root at a distance to the RAM (Fig. 1b,g). In this region, also *OPS* and *OPL2* promoters displayed the expected activity in the newly developing phloem cells (Fig. 1f,h) (Ruiz-Sola *et al*., 2017). Conversely, *OPL3* and *OPL4* promoters were inactive in the more developed part of the root (Fig. 1i,j).

**Figure 1:**
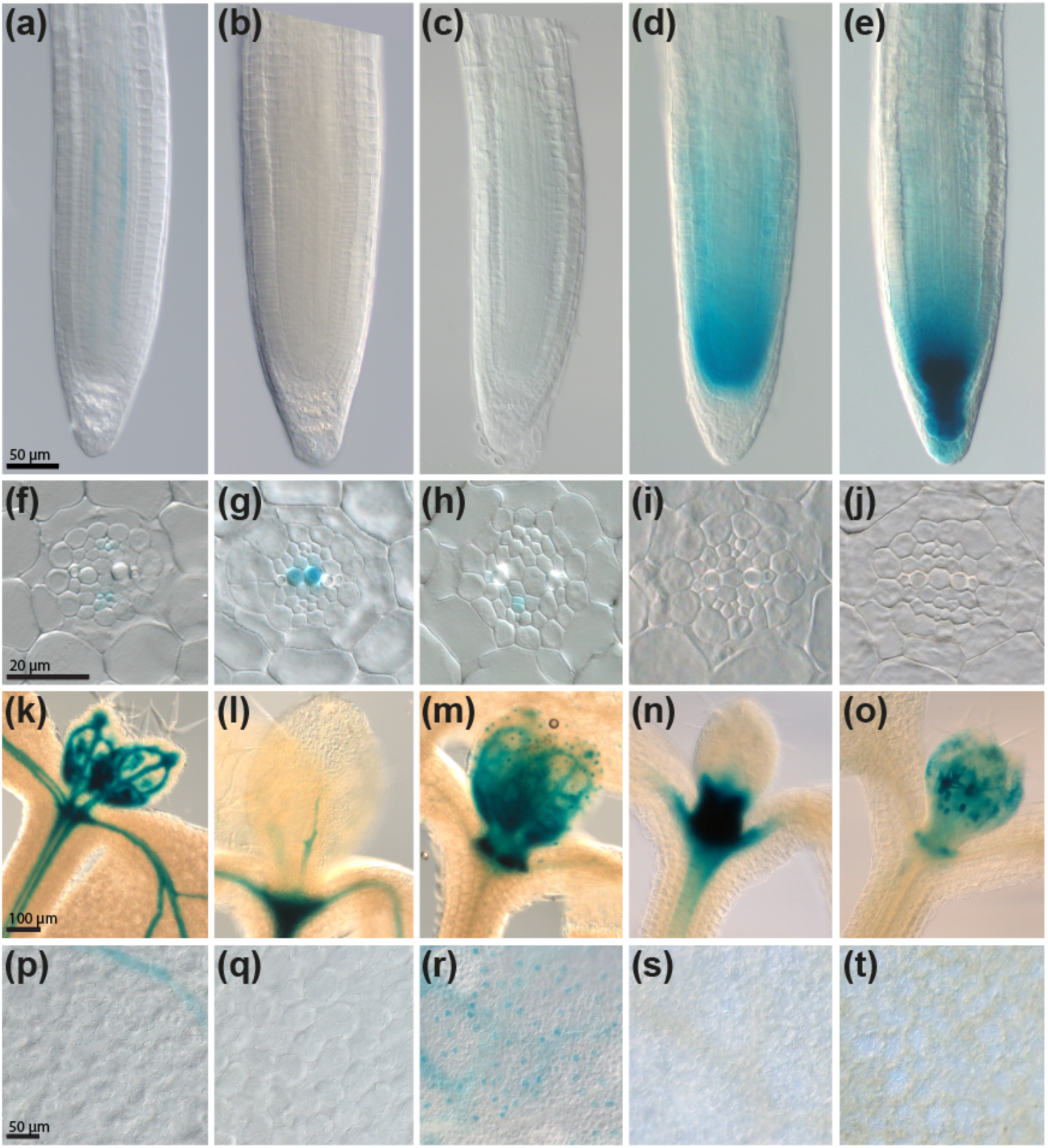
Promoter activity of all *OPS/OPL* family genes. Promoter activities of *OPS/OPL* family genes were monitored in 5-day-old Arabidopsis seedlings carrying promoter*-GUS* constructs. Shown is GUS activity in **(a - e)** root tips, **(f - j)** sections through the root stele in the early differentiation zone, **(k - o)** first emerging true leaves and shoot meristem, and **(p - t)** cotyledon epidermis. *OPS* promoter activity is depicted in **(a, f, k, p)**, *OPL1* promoter activity in **(b, g, l, q)**, *OPL2* promoter activity in **(c, h, m, r)**, *OPL3* promoter activity in **(d, i, n, s)**, and *OPL4* promoter activity in **(e, j, o, t)**. Scale bar information is included in the images, and images representing the same tissue type are consistently scaled.

In the seedling shoot, all class I genes (*OPS*, *OPL1*, *OPL2*) displayed vascular expression (Fig. 1k,l,m). In *OPL3* promoter-*GUS* plants, strong GUS activity was detected in stipules and extended slightly towards the vasculature immediately adjacent to the shoot apical meristem (Fig. 1n). The *OPL4* promoter was active in the epidermis of young leaves, mainly in developing trichomes (Fig. 1o), but not in mature trichomes (Fig. 1o,t). Moreover, we observed GUS activity in the developing guard cells of *OPL2* reporter plants (Fig. 1r), but not in those of *OPS*, *OPL1*, *OPL3*, or *OPL4* reporter plants (Fig. 1p,q,s,t).

To compare our results for promoter activity with OPS/OPL protein localisation, we generated and investigated lines expressing fusions of the *OPS*/*OPL* genes with *GREEN FLUORESCENT PROTEIN* (*GFP*) driven by the same promoters used for the GUS-reporter plants. Fusing GFP to the N-terminus of OPS and OPL2 was previously shown not to inhibit the function of these proteins (Ruiz-Sola *et al*., 2017). As expected, in 5-day-old roots, OPS-GFP was exclusively located in the protophloem and incipient metaphloem cell files, starting from the phloem initial (Fig. 2a,a’,b) (Truernit *et al*., 2012; Rodriguez-Villalon *et al*., 2014). While the OPL1-GFP fusion protein was only observed in developing metaxylem cells, and not in the RAM (Fig. 2c,c’,d, Fig. S1), OPL2-, OPL3-, and OPL4-GFP were clearly detectable in the RAM. Again, fluorescence in OPL4-GFP plants was slightly stronger around the quiescent centre of the root and extended towards the columella cells (Fig. 2i,i’) while OPL3-GFP fluorescence did not (Fig. 2g,g’). Overall, the patterns of protein localisation in the RAM corresponded well with our promoter activity data. Fluorescence of OPL3- and OPL4-GFP in the RAM was also again notably stronger than that of OPL2-GFP, as indicated by the need to increase the laser-power of the confocal laser-scanning microscope when imaging the latter (Fig. 2e,e’).

**Figure 2:**
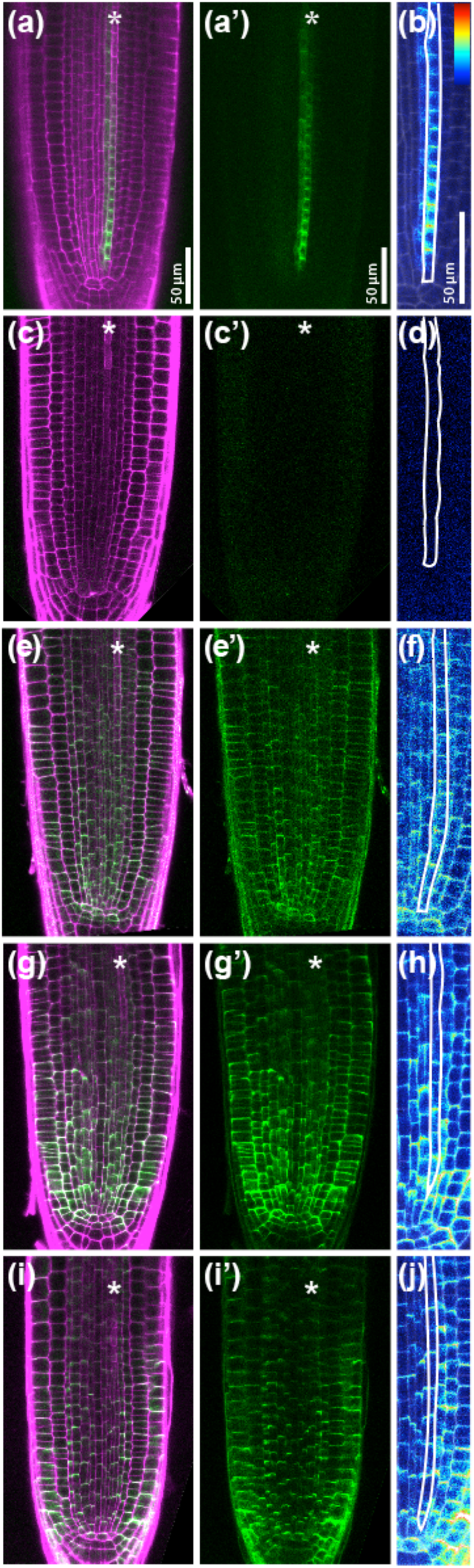
Localisation of OPS/OPL-GFP fusion proteins in root tips. Confocal laser-scanning images of 5-day-old Arabidopsis root tips expressing *OPS-GFP* **(a, a’, b)**, *OPL1-GFP* **(c, c’, d)**, *OPL2-GFP* **(e, e’, f)**, *OPL3-GFP* **(g, g’, h)**, and *OPL4-GFP* **(i, i’, j)** under control of their native promoters. **(a, c, e, g, i)** show overlay images of GFP fluorescence (green) and propidium iodide staining of cell outlines (magenta), while **(a’, c’, e’, g’, i’)** display the GFP channel only. The protophloem cell file is marked with an asterisk. **(b, d, f, h, j)**. Localisation of the fusion proteins in the protophloem region (outlined in white), with GFP fluorescence colour-coded for intensity levels (see insert in **(b)**, with red representing high and blue representing low intensity). Scale bar information is included in the images, and images representing the same tissue type are consistently scaled.

In summary, OPL2, OPL3, and OPL4 exhibited broad and overlapping localisation in the RAM with slight differences in their overall and tissue specific expression strengths. In contrast, OPS and OPL1 displayed narrow expression patterns in the developing phloem and metaxylem cell files, respectively. In leaves, only class I genes were expressed in the vasculature.

### OPS/OPL proteins are polarly localised and differentially expressed in the protophloem cell files

Our protein localisation data confirms previous findings that in their root expression domains all OPS/OPL proteins are polarly localised to the basal (shootward) part of the cell (Fig. 2) (Truernit *et al*., 2012; Ruiz-Sola *et al*., 2017; Wallner *et al*., 2023). While not the focus of this work, we noted that the GFP-fusion proteins were often more abundant in established cell walls, with less protein associated with the newly formed cell walls after cell divisions (e.g. Fig. 2f).

To understand the contribution of all OPS/OPLs to phloem differentiation, we further investigated their expression in the developing root protophloem files. While OPS-GFP was exclusively localised in these cell files, OPL2-, OPL3-, and OPL4-GFP fusions were all also clearly present in this tissue. Although, due to the polar localisation of the GFP-fusions, quantifying relative expression strength is challenging, it appeared that OPL2 expression was slightly stronger, whereas OPL3 and OPL4 displayed slightly weaker expression in the developing protophloem when compared to their expression in the surrounding cell files (Fig. 2f,h,j). Regardless of this, it can be concluded that, except for OPL1, all OPS/OPL proteins are expressed and polarly localised in the developing protophloem sieve tube cell files.

### OPLs exhibit functional overlap with OPS

Expressing *OPL2-GFP* under the control of the *OPS* promoter in *ops* can alleviate OPS loss-of-function phenotypes (Ruiz-Sola *et al*., 2017). Here, we replicated this experiment for OPL1, OPL3, and OPL4. Fig. 3a illustrates that two of three representative single-insert homozygous T3 lines selected for *proOPS-OPL1-GFP* and *proOPS-OPL3-GFP* each fully restored the *ops* short root phenotype. Of the three selected *proOPS-OPL4-GFP* lines, one line exhibited significant rescue of *ops* root growth (Fig. 3a). Moreover, all complementation lines exhibited phloem-specific expression and OPS-like polar localisation of the OPL-GFP fusion proteins (Fig. S2a-e).

**Figure 3:**
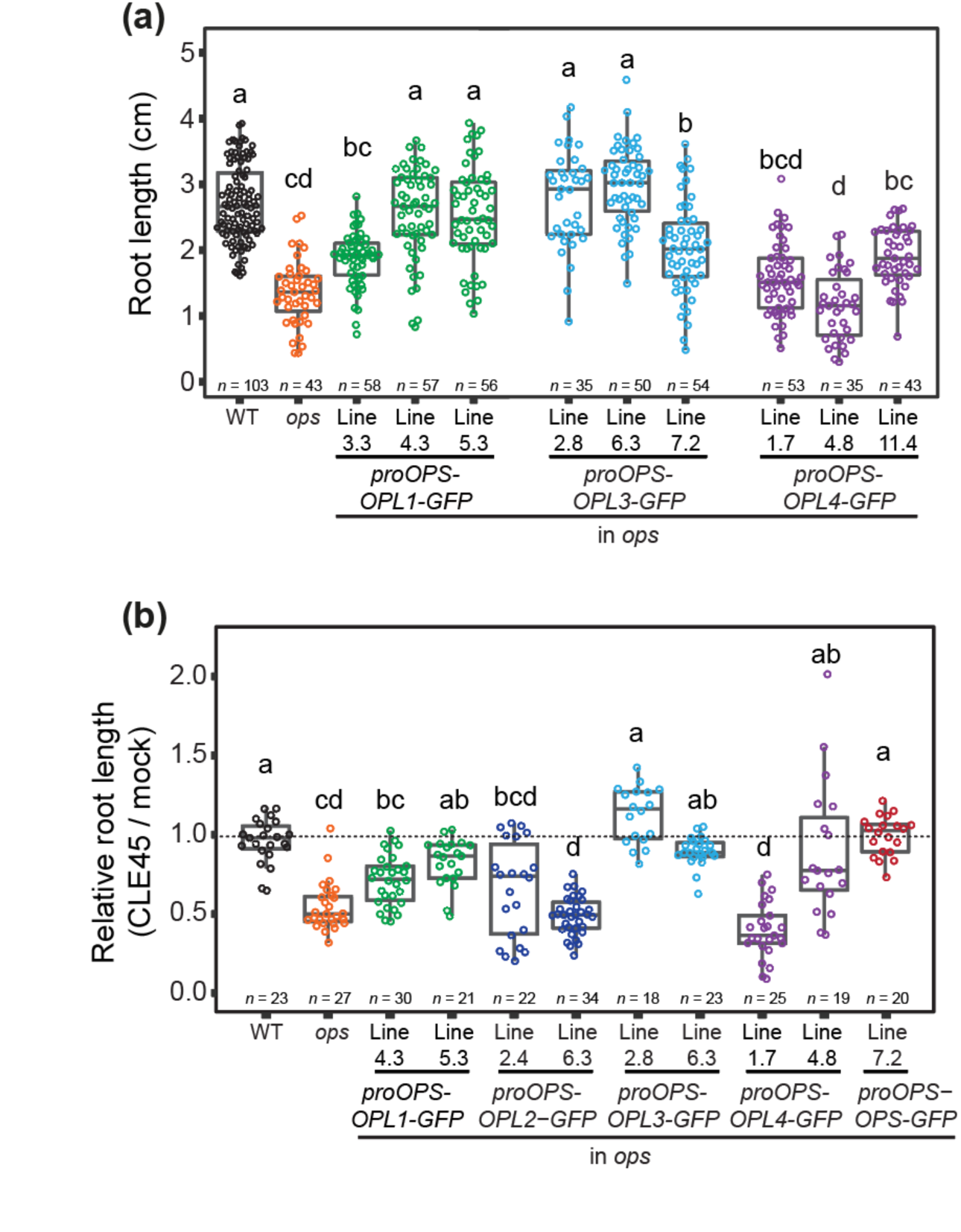
Root growth of *ops* plants expressing *OPS/OPL-GFP* fusion proteins under control of the *OPS* promoter on normal and CLE45 supplemented media. (**a**) Root length of 8-day-old *ops* seedlings harbouring the specified constructs, with three independent homozygous single-insert lines displayed for each. **(b)** Root length of selected lines germinated and grown for 6 days on media supplemented with 5 nM CLE45 relative to their respective length on control media. *n* represents the number of roots analysed. Statistical significance was assessed using a Kruskal-Wallis test followed by Dunn’s multiple comparison post hoc test and Bonferroni *p* value adjustment. Different letters indicate statistically significant differences (*p* < 0.05). Wild type (WT) and *ops* serve as controls.

We also assessed one of the best complementing lines for each gene for complementation of the *ops* vascular complexity phenotypes. Overall, improvements of vascular complexity was observed in all lines. The line expressing *OPL1-GFP* under the control of the *OPS* promoter in *ops* showed the weakest rescue, while the expression of *OPL3-GFP* in this context provided a rescue similar to that of *proOPS-OPS-GFP* in *ops* (Fig. S2f,g).

Consistent with a role of OPS in attenuating CLE45 signalling in the protophloem (Rodriguez-Villalon *et al*., 2014; Breda *et al*., 2019; Carbonnel *et al*., 2023), *ops* root growth was more susceptible to low CLE45 concentrations (Fig. 3b). Since most of our complementation lines exhibited better root growth than *ops*, we also investigated whether CLE45 sensitivity was restored to wild-type-levels in these lines. For this, we chose two well-complementing lines for each OPL complementation construct and a CLE45 concentration which only inhibited *ops* but not wild type root growth. In this assay, the best root growth complementing lines also exhibited a CLE45 tolerance similar to the wild type, while for the complementation lines displaying only a partial root growth rescue still a trend towards better relative growth on CLE45 supplemented media compared to *ops* was observed (Fig. 3b).

In summary, all OPL proteins can partially compensate for OPS loss-of-function and interfere with CLE45 signalling when expressed in the *OPS* expression domain. This implies that, to some extent, all OPL proteins are functionally equivalent to OPS. While we observed quantitative differences in the ability of individual *OPL* genes to rescue specific aspects of the *ops* mutant phenotypes, currently, we cannot distinguish whether these differences reflect variations in the protein’s functionality or slight variations in expression strength or quality due to positional effects resulting from the genomic insertion site of the complementation constructs.

### Single and multiple mutant combinations reveal a major role for OPS in plant growth control

While *opl2* root growth is wild-type-like, *ops* roots grow more slowly, and their meristem size and root diameter are reduced (Truernit *et al*., 2012; Rodriguez-Villalon *et al*., 2014; Ruiz-Sola *et al*., 2017). Here, we examined T-DNA insertion lines for *OPL1, OPL3,* and *OPL4,* which were in part different to the ones used by Wallner et al (Fig. S3a,b) (Alonso *et al*., 2003; Wallner *et al*., 2023). Although RT-PCR confirmed that there was no residual transcript in these lines (Fig. S3c-e), we did not observe any mutant root phenotypes (Fig. S4a-c), including in the metaxylem of *opl1* roots (Fig. S4d,e). Also the trichomes of *opl4* plants were wild-type-like (Fig. S4f,g). Thus, among the *ops/opl* single mutants, only *ops* exhibits a mutant phenotype in standard growth conditions.

Overlap of expression patterns and functional interchangeability, however, suggested redundancy among *OPS/OPL* genes. To test this, we generated several higher-order mutant combinations. First, we generated plants in which all *OPL* genes broadly expressed in the root meristem (*OPL2*, *OPL3*, *OPL4*) were knocked out. Root growth, root diameter, and meristem size of *opl2 opl3 opl4* plants, however, were wild-type-like, and an additional knock-out of *OPL1* in this background did not change this (Fig. S4h-j). We also tested CLE45 sensitivity of *opl1 opl2 opl3 opl4*. Unlike *ops*, however, *opl1 opl2 opl3 opl4* was not hypersensitive to CLE45, although three of the four OPS/OPL proteins present in the developing protophloem and capable of interfering with CLE45 signalling were absent in this mutant (Fig. S4k).

Next, we produced mutant combinations with *ops*. The loss-of-function of individual *OPL* genes in the *ops* background led to *ops-*like phenotypes for all combinations, except *ops opl2*, which, as previously shown, displayed significantly more impaired root growth than *ops* (Fig. 4a-c) (Ruiz-Sola *et al*., 2017). Fig. 4 illustrates a trend found in some of our assays towards slightly reduced root growth for certain *opl* mutant combinations with *ops*. However, this was inconsistent across experiments, at best suggesting a higher sensitivity of these combinations to minor variations in the experimental conditions.

**Figure 4:**
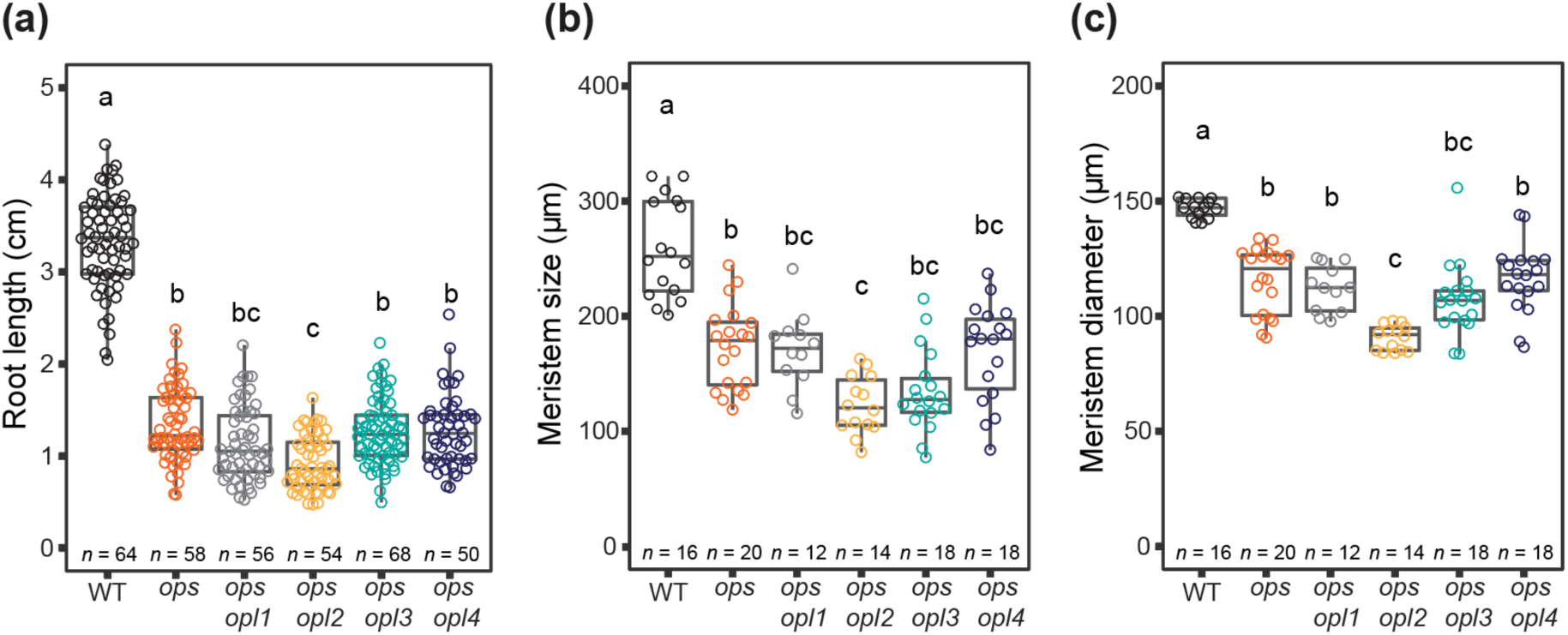
Root phenotypes of *ops opl* double mutant combinations. **(a)** Primary root length of 8-day-old seedlings. **(b)** Root meristem length and **(c)** diameter of 5-day-old seedlings. *n* represents the number of roots analysed. Statistical significance was assessed using a Kruskal-Wallis test followed by Dunn’s multiple comparison post hoc test and Bonferroni *p* value adjustment. Different letters indicate statistically significant differences (*p* < 0.05). Wild type (WT) and *ops* serve as controls.

### A knockout of all *OPS/OPL* genes uncovers a high degree of redundancy within the gene family

Redundancy within the *OPS/OPL* gene family became fully apparent when we generated the quintuple mutant *ops opl1 opl2 opl3 opl4* (hereafter called *opx qm*). Confirming the findings of Wallner and colleagues, shoot growth was significantly impaired in *opx qm* (Fig. 5) (Wallner *et al*., 2023). Compared to *ops opl2*, which already shows reduced shoot growth (Fig. 5a,b), *opx qm* rosettes were even smaller and grew more slowly, with delayed flowering, decreased seed production, and failure to reach the height of *ops opl2* (Fig. 5a-c). The length of cortex cells in the fully elongated parts of the flowering stems of these plants, however, was similar in *opx qm* and *ops opl2* (Fig 5d). Conversely, vascular complexity was further reduced in *opx qm*, also when compared to *ops opl1 opl2*, in which all OPS/OPL proteins present in the leaf vasculature are missing (Fig. 5e,f).

**Figure 5:**
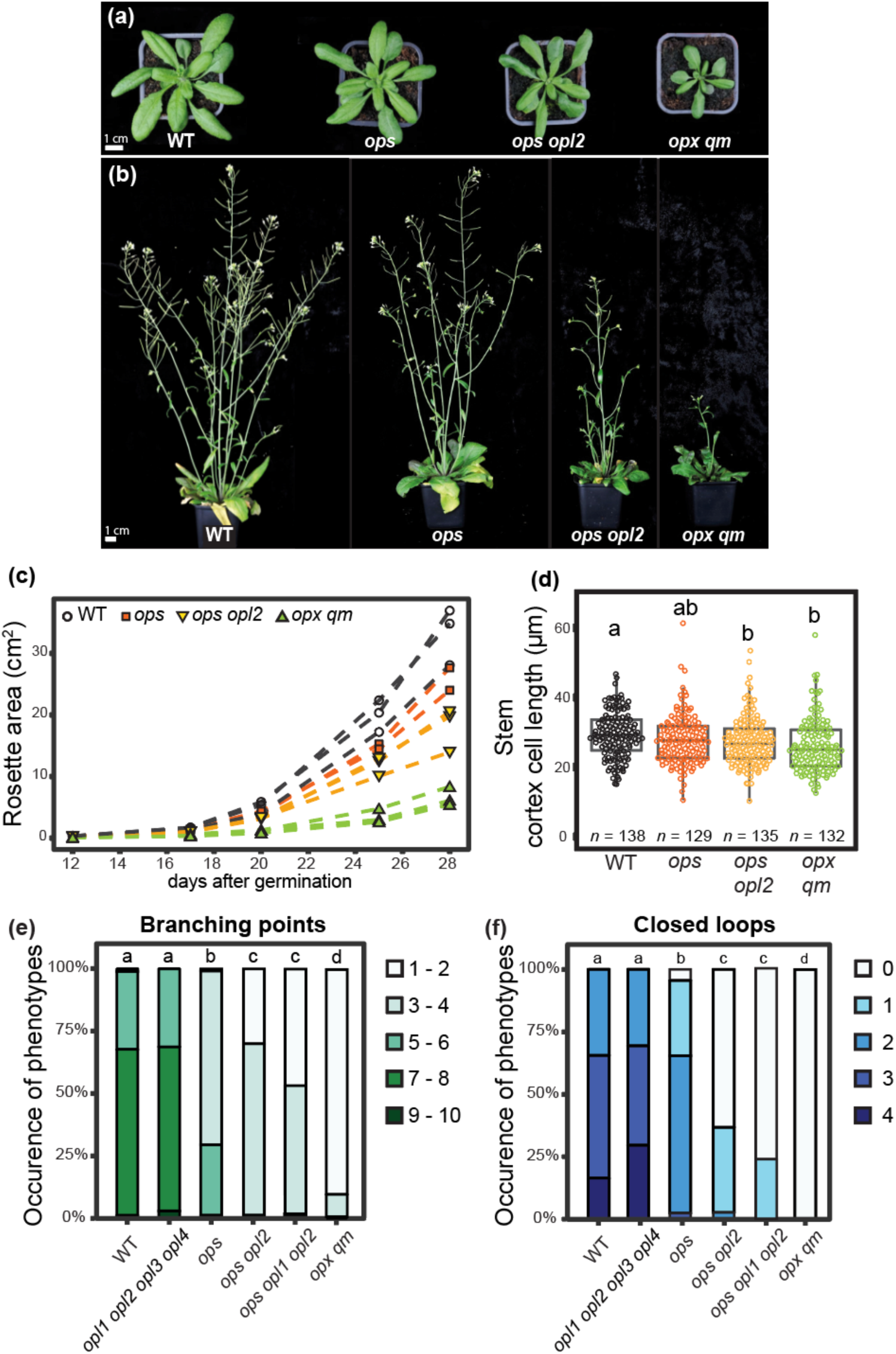
Shoot phenotypes of *ops/opl* higher-order mutants. Shoot phenotypes representative of the indicated genotypes. **(a)** Rosettes of 28-day-old plants. (**b**) Shoots of 42-day-old plants. **(c)** Rosette leaf area measurements of individual plants over several weeks. Scale bar information is included in the images. **(d)** Length of cortex cells in fully elongated stem parts. Five plants were analysed for each genotype, *n* represents the number of cells analysed. Statistical significance was assessed using a Kruskal-Wallis test followed by Dunn’s multiple comparison post hoc test and Bonferroni *p* value adjustment. **(e, f)** Vascular complexity phenotypes of cotyledons of 7-day-old seedlings. For each genotype, 90 to 130 cotyledons were analysed. Statistical significance of cotyledon phenotypes was calculated by Fisher’s exact test on the 5 vascular categories indicated. Different letters indicate statistically significant differences (*p* < 0.05). Wild type (WT) and *ops* serve as controls.

In roots, the meristem size was more reduced in *opx qm* than in *ops opl2*, while the diameter of the meristem was similar to *ops opl2* (Fig. 6a-c, see also Fig.7a-d). There was also no difference between the two genotypes in the length of cortex cells in the meristem, but, consistent with the reduced meristem size, cells started elongating earlier (Fig. 6d). Expectedly, root length was also further reduced in *opx qm* (Fig. 6a,e). Despite being visually apparent earlier (Fig. 6a), this became statistically significant only after longer growth times, possibly due to higher measurement error when assessing the very short roots of younger plants (Fig. 6e). To measure the root length of older plants, we had to switch to a 12 hours light / 12 hours dark growth cycle. This change was necessary because only under these conditions could we still distinguish between the primary and lateral or adventitious roots in our mutants (Fig. S5a,b).

**Figure 6:**
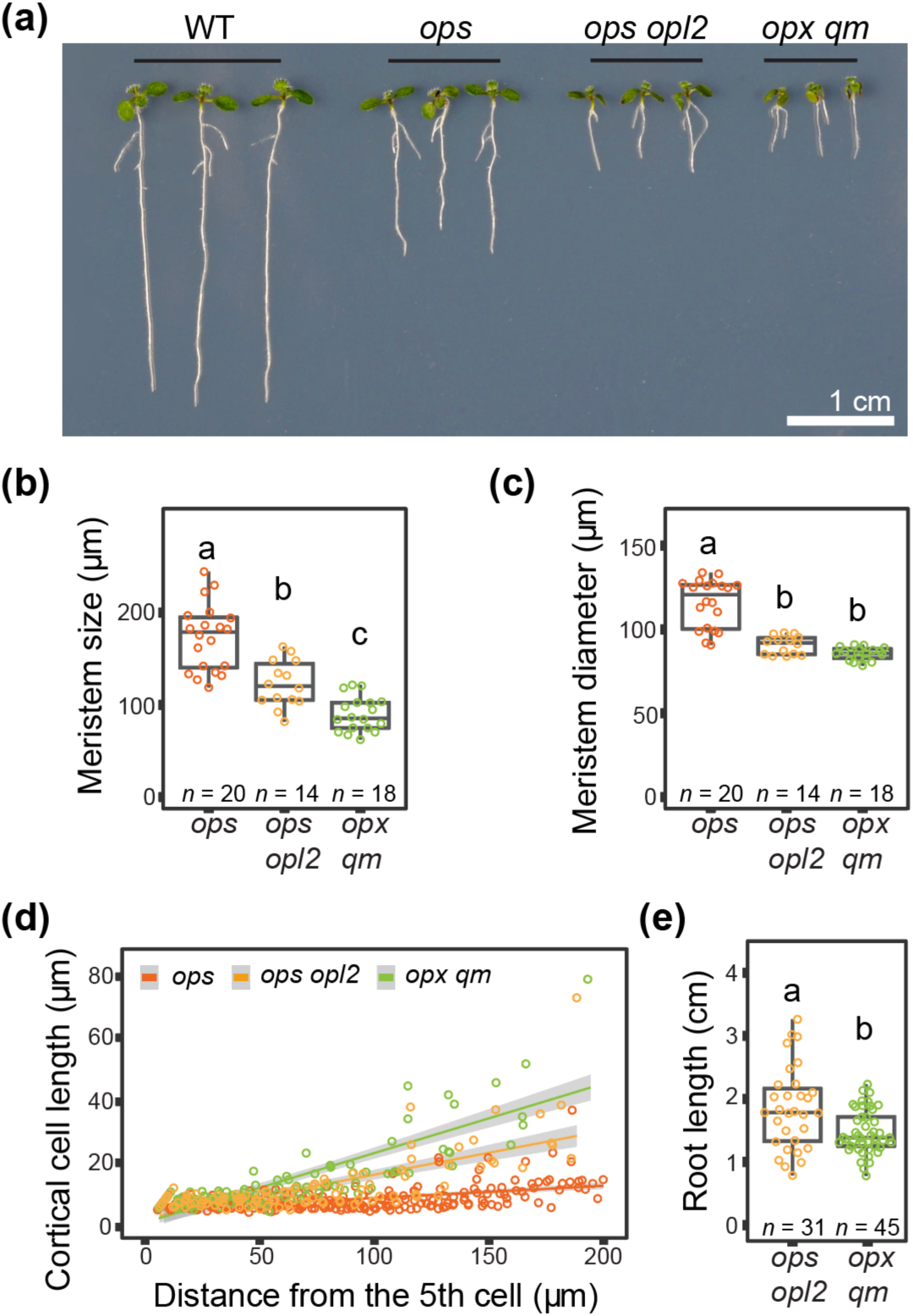
Root phenotypes of *opx qm* compared to *ops* and *ops opl2*. **(a)** Root phenotype of 7-day-old plants of the indicated genotypes. The wild type (WT) is included for comparison. Scale bar information is included in the image. **(b)** Root meristem size and **(c)** diameter of 5-day-old seedlings. **(d)** Length of root cortex cells of 5-day-old roots relative to their position in the meristem (position of cell relative to the 5th cell counting from the QC onwards). The line represents the linear trend, the 95% confidence interval is depicted in grey. **(e)** Primary root length of 14-day-old seedlings grown under a 12/12-hour light/dark regime. *ops* was not included in this comparison. *n* represents the number of roots analysed. Statistical significance was assessed using a Kruskal-Wallis test followed by Dunn’s multiple comparison post hoc test and Bonferroni *p* value adjustment. Different letters indicate statistically significant differences (*p* < 0.05).

The presence of more severe phloem defects would likely have further reduced the overall growth of *opx qm* compared to *ops opl2*. However, in transport assays, a similar number of plants in both genotypes failed to transport the mobile dye 5(6)carboxyfluorescein diacetate (CFDA) from the shoot to the root tip (*ops opl2*: 3/30, *opx qm*: 3/29, WT: 0/20). Confocal laser-scanning images of the root protophloem and root cross-sections further supported the notion that the phloem defects in *opx qm* and *ops opl2* were similar, as there was no increase in the number of undifferentiated cells in the proto- and metaphloem sieve tube positions in *opx qm* compared to *ops opl2* (Fig. 7a-j). Additionally, the total number of cells in the stele was also similar in the two genotypes (Fig. 7k).

**Figure 7:**
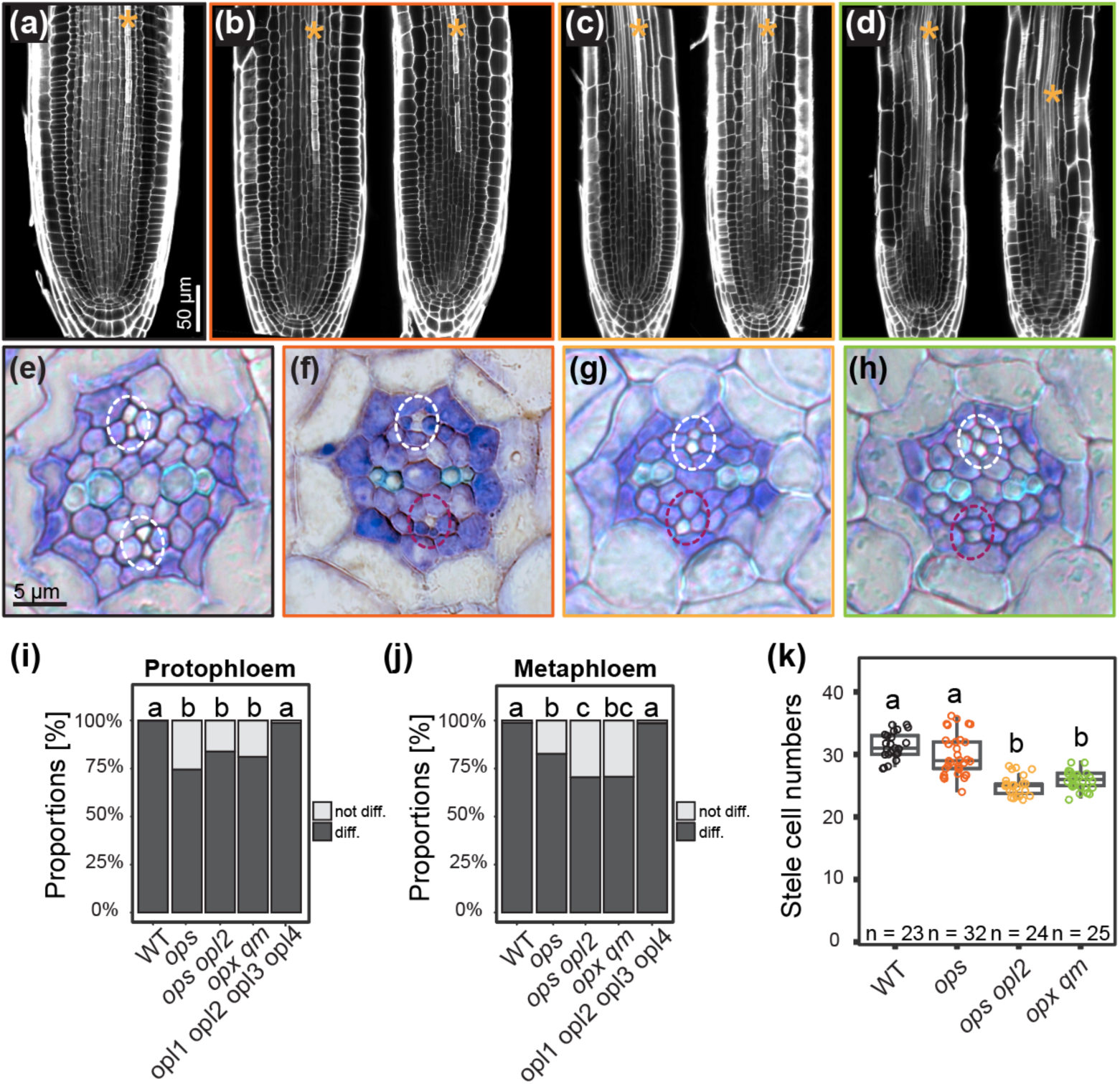
Stele phenotypes of *ops/opl* higher-order mutants. **(a - d)** Confocal laser-scanning images of root meristem cellular architecture of 5-day-old roots, with representative examples for **(a)** wild type, **(b)** *ops*, **(c)** *ops opl2*, and **(d)** *opx qm*. For each mutant, an example of a root displaying an intact (left) and an interrupted (right) protophloem file (marked with an asterisk) is shown. **(e - h)** Examples of toluidine blue stained stele cross sections of 6-day-old roots with representative phenotypes of **(e)** wild type, **(f)** *ops*, **(g)** *ops opl2*, and **(h)** *opx qm*. The phloem poles are encircled (white, if wild-type-like, dark red if deviating from typical wild-type phenotype). Cells appearing white are fully differentiated sieve tubes. Scale bar information is included in the images, and images representing the same tissue type are consistently scaled. **(i, j)** Quantification of the relative number of undifferentiated **(i)** protophloem and **(j)** metaphloem sieve tube cells in cross sections. For each genotype, 80 to 100 phloem poles were analysed. Statistical significance of phloem phenotypes was calculated by Fisher’s exact test. **(k)** Number of cells in the stele of the indicated genotypes. *n* represents the number of roots analysed. Statistical significance was assessed using a Kruskal-Wallis test followed by Dunn’s multiple comparison post hoc test and Bonferroni *p* value adjustment. Different letters indicate statistically significant differences (*p* < 0.05).

Overall, these data suggest that, besides the role of *OPS* and *OPL2* in phloem development, *OPS/OPL* genes together control plant growth vigour, likely by positively influencing meristematic activity in plants. Further, OPS is not differently expressed in *opl2 opl3 opl4*, ruling out the possibility that OPS takes over the function of the OPL proteins when they are not expressed (Fig. S5c).

### Brassinosteroid growth response differs in *ops* and *opx qm*

Reduced organ growth, cell size, and meristem cell number are typical features of mutants defective in BR biosynthesis or signalling. Given that OPS can negatively regulate BIN2 activity and BIN2 is broadly expressed in the root meristem (Anne *et al*., 2015), a potential cause for the observed *opx qm* growth phenotypes could be the involvement of all OPS/OPL proteins in modulating the BR signalling pathway by inhibiting BIN2 within their expression domains. However, the root growth of *opl2 opl3 opl4* (absence of all OPLs broadly expressed in the root meristem) on bikinin (a BIN2 inhibitor), brassinolide (a brassinosteroid), and brassinazole (a BR biosynthesis inhibitor) was wild-type-like (Fig. 8a-c). Conversely, in line with BIN2 hyperactivity in *ops*, the root growth of *ops* clearly improved when grown on bikinin or brassinolide, whereas *opx qm* root growth showed less improvement when transferred to media supplemented with these compounds (Fig 8a,b). These findings suggest that the reduced root growth observed in *opx qm* is not merely a consequence of BIN2 hyperactivity in this line.

**Figure 8:**
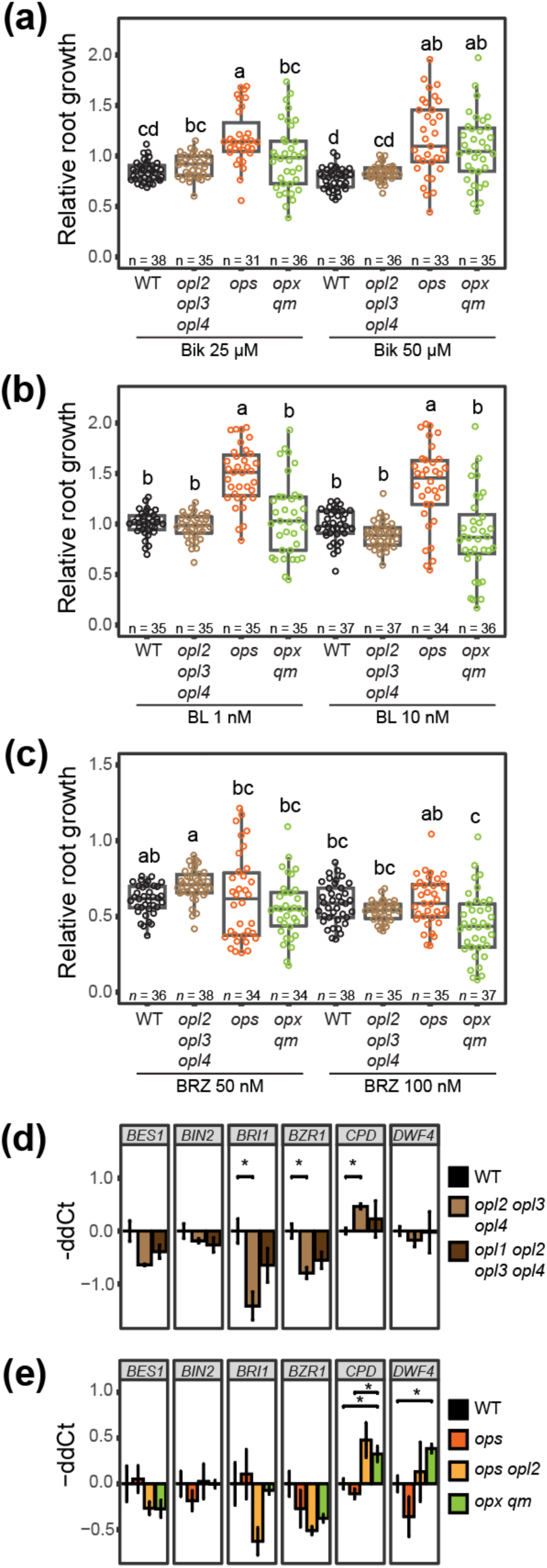
BR related phenotypes of *ops/opl* higher-order mutants. **(a - c)** Relative root growth of the indicated genotypes after transfer to media containing the indicated amount of **(a)** bikinin (Bik), **(b)** brassinolide (BL), and **(c)** brassinazole (BRZ). Growth is relative to the respective growth on control media. *n* represents the number of roots analysed. Statistical significance was assessed using a Kruskal-Wallis test followed by Dunn’s multiple comparison post hoc test and Bonferroni *p* value adjustment. Different letters indicate statistically significant differences (*p* < 0.05). Wild type (WT) and *ops* serve as controls. **(d, e)** Quantification of BR related gene expression in the indicated genotypes by qPCR. The results represent the means ± SEM of three biological replicates. A ddCt of 1 represents a two-fold increase, whereas a ddCt of -1 represents a 0.5-fold decrease in expression compared to the wild-type. Stars above the bars indicate significant differences (*p* < 0.05) as determined by unpaired two-tailed Student’s t-test.

Nevertheless, qPCR analysis in 8-day-old seedlings revealed upregulation of BR biosynthesis enzyme gene expression (*DWF4*, *CPD*) in several *ops/opl* mutant combinations, including those without *ops* (Fig. 8d,e). This finding would point more towards an overall downregulation of BR signalling in these lines. Indeed, *BRI1* and *BZR1* expression were significantly downregulated in *opl2 opl3 opl4*, with a tendency towards downregulation of expression of these genes also in higher-order mutant combinations with *ops* (Fig. 8d,e). Although caution is needed in interpreting these experiments performed at the whole plant level, they nevertheless indicate an imbalance in BR signalling pathway components in *opl* multiple mutant combinations with or without *ops*.

## Discussion

In this study, we conducted a comprehensive analysis of the Arabidopsis *OPS/OPL* gene family, focusing on root and vascular development. We show that in leaves, only class I gene expression is linked to the vasculature. In roots, two of the three class I genes, *OPS* and *OPL1* exhibit a vascular specific expression pattern in phloem and xylem cells, respectively. While *OPL2* is broadly expressed in the root meristem (Ruiz-Sola *et al*., 2017), judging from OPL2-GFP fluorescence, its expression appears to be stronger in the developing phloem files in the root. In the differentiated part of the root, *OPL2* expression it is specific to the phloem. Similarly to *OPL2*, the class II genes *OPL3* and *OPL4* are broadly expressed in the root meristem, but their expression is relatively weaker in the developing root phloem. Apart from *OPL1* expression, which was reported by others to be expressed in the root meristem, our data align well with previous findings (Nagawa *et al*., 2006; Wallner *et al*., 2023). For *OPL2*, our data also confirm that its promoter is strongly active during guard cell development, while we may have overlooked the reportedly weak promoter activity for *OPL1*, *OPL3*, and *OPL4* in the guard cells of our promoter-*GUS* plants (Wallner *et al*., 2023). Interestingly, the *OPL2* promoter-driven GUS activity in the root meristem was very weak, while we clearly observed expression of the OPL2-GFP fusion protein under control of the same promoter. This suggests that OPL2 could be a rather stable protein.

Despite their different expression patterns, when expressed under control of the *OPS* promoter in *ops,* all OPL proteins can assume the function of OPS, including the dampening of CLE45 signalling. While there may be quantitative differences in functional equivalence among the proteins, this finding suggests that their molecular function is similar. However, as CLE45 is exclusively expressed in the developing phloem (Rodriguez-Villalon *et al*., 2014), the function of *OPL* genes almost certainly extends beyond interfering with CLE45 signalling, potentially involving other pathways associated with receptor kinase signalling. In essence, this strongly indicates that the specific expression domains of the OPS/OPL proteins are pivotal for determining their functions.

For our analysis, we generated several multiple mutant combinations, including the quintuple mutant *opx qm*, which was also independently produced by Wallner and colleagues, who observed similar shoot phenotypes using different alleles (with T-DNA insertion sites further downstream) for *opl3* and *opl4* (Wallner *et al*., 2023). Since the growth of *opl1 opl2 opl3 opl4* was wild-type-like while *opx qm* growth was severely impaired, our results generally emphasise the central role for OPS in plant growth.

In terms of phloem development, *opx qm* did not exhibit more severe defects than *ops opl2* when assessing the number of undifferentiated sieve elements in the proto- and metaphloem position. Additionally, unlike *ops*, *opl2 opl3 opl4* did not show hypersensitivity to CLE45 treatment. Given that OPL2, OPL3, and OPL4 are all present in the developing phloem and our *ops* complementation assays show that they are capable of interfering with CLE45 signalling in the phloem domain, this result was rather surprising. While, in this respect, testing CLE45 sensitivity of *ops opl2* or *opx qm* would have been interesting, it was technically not feasible due to their already significantly reduced root growth. Overall, the most plausible explanation for these results is twofold. First, among the OPS/OPL proteins, OPS and OPL2 could be best adapted to promote phloem sieve tube differentiation. Second, when considering the expression strength of the *OPS/OPL* genes in the phloem and the rest of the RAM, ensuring a relatively higher dosage of OPS/OPL protein within the phloem domain compared to the surrounding tissues may actually be the crucial factor for proper phloem development. This idea is also supported by our *ops* complementation assays.

While there was no redundancy in phloem development, the shoot and root growth phenotypes clearly indicated redundant functions of OPL1, OPL3, and OPL4 with OPS and OPL2. *Opx qm* exhibited significantly more impaired shoot and root growth than *ops opl2*. Since *ops opl2* roots were already very short, disentangling the individual contributions of OPL1, OPL3, and OPL4 to root growth would have been too challenging. Their contributions to shoot growth, however, were more apparent. Regarding vascular complexity, we were able to demonstrate that an additional knock-out of *OPL3* and *OPL4* in the class I mutant further reduced vascular complexity, indicating an additive role for the class II genes in this process. The contribution of OPL1 to plant growth is the least clear from our experiments.

The absence of more severe phloem defects in *opx qm*, as judged from CFDA transport data and phloem anatomy, rules out the possibility that the reduction in growth compared to *ops opl2* is solely due to more severe solute transport problems in *opx qm*. The heavily impaired growth of *opx qm*, coupled with the fact that we did not find reduced cell sizes in *opx qm* when compared to *ops opl2*, strongly suggests a role of the *OPL* genes in meristem activity. A recurring theme in OPS/OPL function appears to be the regulation of the transition from cell division to differentiation. While the phenotypes observed in *ops* mutants and mild overexpressors align with a role of OPS in promoting protophloem differentiation (Ruiz-Sola *et al*., 2017; Breda *et al*., 2019), OPL2 has been suggested to promote stemness during guard cell development and also when heterologously expressed in Marchantia (Wallner *et al*., 2023). A role in maintaining cell division would also be consistent with the presence of OPL2, OPL3, and OPL4 specifically in the root meristem. Furthermore, an imbalance in the transition from cell division to cell elongation in distinct tissues could well lead to the observed general growth retardation in the mutants, with an imbalance of these processes in the phloem, as a central organiser of plant growth, having the worst effect.

Although *opx qm* root meristems only partially resemble those of classical BR mutants (Kang *et al*., 2017; Vukašinović *et al*., 2021; Fridman *et al*., 2021) - for example, we found cell length in the meristem to be unaffected - the number of cells in the meristem is clearly reduced. Moreover, the number of stele cells is also reduced in *opx qm* compared to *ops*, which could also indicate altered BR signalling (Kang *et al*., 2017). In this regard, it was interesting to observe that *ops*, *opl2 opl3 opl4*, and *opx qm* reacted differently to treatments with bikinin or brassinolide. The enhanced growth of *ops* on bikinin or brassinolide, which was reported before (Anne *et al*., 2015; Kang *et al*., 2017), is consistent with a role of OPS in inhibiting BIN2, thereby promoting BR response. The relatively reduced reaction of *opx qm* and the wild-type-like growth behaviour of *opl2 opl3 opl4* on bikinin and brassinolide could be explained with a role of OPL2, OPL3, and OPL4 in dampening this BR response. Alternatively, our observations, especially the fact that we see upregulation of *DWF4* and *CPD* expression in several mutant combinations, would also align with a role of the OPLs in promoting BR biosynthesis by inhibiting other GSK3-like kinase in Arabidopsis, some of which are not sensitive to bikinin (De Rybel et al., 2009). Due to the complex and concentration dependent nature of BR action (Vukašinović *et al*., 2021) and the distinct but overlapping localisation of OPS/OPL proteins, more spatially resolved data will be required to support this notion.

Taken together, our data demonstrate that among the *OPS/OPL* gene family, only *OPS* and *OPL2* play a crucial role in the development of continuous phloem sieve tubes. Furthermore, our findings suggest a general role of the *OPS/OPL* genes in enhancing plant growth vigour by modulating and possibly coordinating cell division and/or differentiation, likely in a context-dependent manner. As membrane-localised proteins, they could either promote or interfere with the interaction of numerous signalling pathway components, including those of the BR and CLE signalling pathways. Further investigations into the role of all *OPL* genes are hindered by the essential role of the phloem in plant growth. Nevertheless, targeted misexpression studies and the identification of interaction partners will provide additional insights into the role of this intriguing gene family in plant development.

## Acknowledgements

We acknowledge the ETH Zurich Scientific Centre for Optical and Electron Microscopy (ScopeM) for providing microscopy facilities and the ETH Zurich Genetic Diversity Centre (GDC) for collaborating on qPCR experiments. We are grateful to Samuel C. Zeeman (ETH Zurich, Switzerland), Markus Geisler (University of Fribourg, Switzerland) and Clara Sanchez-Rodriguez (CBGP Madrid, Spain) for helpful discussions. Additionally, we want to thank Andrea Ruckle for her assistance with plant work, and Bianka Horváth, Olivia Bärtschi, and Melina Wüstner for technical support during their training periods in our lab. This work was funded by ETH Zurich Research Grants (ETH-33 12-1, ETH-26 17-2). K.B. is supported by a research grant from the Swiss National Foundation (310030_184762).

## Competing interests

We have no competing interests to declare.

## Author contributions

S.C., M.A.R.S., K.B., M.C. and E.T. designed and performed experiments and analysed data. S.S.K.H. performed experiments and analysed data. S.C. and E.T. wrote the manuscript. E.T. conceived and directed the research.

## Notes

### Competing Interest Statement

The authors have declared no competing interest.

